# *Bacillus subtilis* ALBA01 can mitigate onion pink root symptoms caused by *Setophoma terrestris*

**DOI:** 10.1101/601633

**Authors:** P Sayago, F Juncosa, A Albarracín Orio, D.F. Luna, G Molina, J Lafi, D.A Ducasse

## Abstract

Soil-borne pathogen *Setophoma terrestris* is the causal agent of pink root of onion, one of the most challenging diseases in onion production. Conventional approaches for managing the disease like solarization, soil fumigation and crop rotation have not been proven effective enough. In this work, we evaluated the biocontrol capacity of *Bacillus subtilis* ALBA01 (BsA01) against *S. terrestris*, in a highly susceptible onion cultivar, both under greenhouse and field conditions. Disease incidence and severity were evaluated together with growth, photosynthesis among other physiological variables and yield parameters. When compared with plants infected with the pathogen, those plants co-inoculated with BsA01 showed significantly less damage and levels of biocontrol above 50%. With regard to physiological parameters, plants challenged with *S terrestris* and inoculated with BsA01 performed as well as the control non-infected plants revealing a growth promotion effect of BsA01 on onion plants.

## Introduction

Onion (*Allium cepa L.)* is an important vegetable crop with a worldwide production of approximately 73 million tons per year, 8% of which is lost because of disease constraints (Rinland et al. 2015). In Argentina, onion production is very important, in terms of quality and quantity, being the main producer country and exporter of dehydrated onions of Latin America (SENASA 2014).

Pink root disease, caused by the soil-borne fungus *Setophoma terrestris*, is one of the most devastating diseases of onion crops under warm environmental conditions (Hansen 1929; de Gruyter et al. 2010). *Setophoma terrestris* is an *Ascomycete* but it is usually considered a member of the *Deuteromycete, Phaeosphaeriaceae* family (Phookamsak et al. 2014). Typical symptoms of pink root include a pinkish coloration of the roots that becomes dark purple with the progress of the disease (Hansen 1929; Lee et al. 2007). Infected plants show diminished water and nutrients uptake, leading to wilting and necrosis (Schwartz and Mohan 2007). These plants also undergo higher demand for assimilates since pathogens manage to manipulate the host metabolism in favor of their own carbohydrate requirement (Berger et al. 2004). Generally, plant-pathogen interaction begins with a series of quick changes causing photosynthetic reduction and an increment of respiration, photorespiration, and invertases enzyme activity (Leal and Lastra 1984; Roitsch et al. 2003). Moreover, a fungal infection may reduce the photosynthetic performance by inhibiting stomatal opening in response to a complex signalling pathway as a way of self-defense (Costa Pinto et al. 2008; Sawinski et al. 2013). Even though onion pink root has been considered as an endemic disease for a long time, not enough research has been conducted to reveal its physiological consequences, especially in regards to the photosynthetic apparatus.

Management of onion pink root is considered a difficult task. Several practices aimed to reduce disease incidence have been applied with variable success. Despite the difficulties, solarization has been slightly effective (Porter et al. 1989) and onion varieties resistant to *S. terrestris* have shown some success (Esfahani et al. 2008; Marzu et al. 2018). Nevertheless, it is not unusual to find *S. terrestris* isolates capable of breaking the resistance of selected cultivars. It has been shown that Argentinean *S. terrestris* isolates are highly virulent (Linardelli et al. 2008a; Linardelli et al. 2008b; Lafi 2011) and break the resistance of onion cultivars. In fact, one of the most popular cultivars grown in Argentina, Val 14, has shown to be fairly sensitive to *S. terrestris* (Lafi 2011).

Rhizosphere microorganisms contribute to plant health by increasing the availability of nutrients and hormonal stimulation inducing tolerance to both biotic and abiotic factors (Singh et al. 2016). In this scenario, the deployment of Plant Growth Promoting Rhizobacteria (PGPR) can be an effective strategy for such purpose (Kloepper and Schroth 1978). This positive characteristic has been exploited by using them as biocontrol agents (BCA). Several *B. subtilis* strains are commercially available as BCA for fungal diseases in crops such as cotton, wheat, rice, chili, etc. (Cai et al. 2017; Xiang et al. 2017; Sajitha et al. 2018). We have previously shown that the strain of *Bacillus subtilis* ALBA01, isolated from the rhizosphere of onion plants, was capable of strongly inhibiting the growth of *S. terrestris in vitro* (Albarracín Orio, et al. 2016a; Albarracín Orio et al. 2016b).

The objective of this work was to evaluate under greenhouse and field conditions, the ability of *Bacillus subtilis* ALBA01 to perform as a BCA against *S. terrestris*.

## Materials and methods

Two trials were carried out, one under greenhouse conditions and the other one under standard field conditions. For both trials, the same materials were used.

### Plant material

The onion cultivar Val 14 was used for both greenhouse and field experiments. We selected this variety of onion since is the most popular cultivar in Argentina and due to its high susceptibility to *S. terrestris* (Lafi 2011).

### Bacterial and fungal strains

*Bacillus subtilis* ALBA01 (BsA01) was used in this study. The bacterium was isolated from onion rhizosphere at the Agricultural Experimental Station INTA Hilario Ascasubi, Buenos Aires Province, Argentina (39 ° 23’57.15 “S, 62 ° 37’30.64” W) (Albarracín Orio et al. 2016a; Albarracín Orio et al. 2016b).

A native strain of *S. terrestris* (St) was used in both assays. St was isolated from Santiago del Estero Province, an onion-producing area of Argentina that is known for presenting the highest incidence and severity of pink root disease than any other onion cropping areas of Argentina (Linardelli et al. 2008a; Linardelli et al. 2008b; Lafi 2011).

### Bacterial inoculum preparation

BsA01 was cultivated in Brain Heart Infusion medium (BHI) agar at 28 °C for 24 h. A single colony was then transferred to BHI broth in a 250 ml flask. After 48 h of incubation at 28 °C and 170 rpm, a bacterial concentration of 3 × 10^8^ cfu ml_-1_ was obtained (1.3 OD) and used for inoculating seeds and plants.

### *S. terrestris* inoculum preparation

*S. terrestris* was inoculated in Potato Dextrose Agar (PDA) in Petri dishes and the culture was kept for 7 days at 25 °C. 50 grams of wheat seeds were placed in 250 ml glass bottles with 25 ml autoclaved distilled water and inoculated with four 5-mm PDA agar plugs containing fungal mycelia. The bottles were shaken daily and kept for 15 days at 25 °C. Later on, the contents of each flask were ground in coffee grinders with 50 ml tap water and finally diluted to 2,5 l (Netzer et al. 1985).

### Greenhouse experiment

#### *Plants inoculation with B. subtillis* ALBA01 *and with the pathogen*

Seeds of onion cv. Val 14 were imbibed in a physiological solution for 24 h at 28 ± 2 °C. Seeds were then sown in speedlings with commercial substrate “Dynamics 2” (http://www.agriservice.com.ar/wp-content/uploads/2014/07/agri-service-dynamics-sustrato-dynamics-2.pdf). After 10 days, when the emergence of the green sprout was visible, half of the seedlings were inoculated with 1 ml of BsA01 culture (OD = 1.3, 3 × 10^8^ cfu ml^-1^).

Thirty days later, plants were transplanted to 3 l pots. Half of the pots were inoculated with *S. terrestris*. The inoculation was performed by mixing vigorously 15 l of the substrate with 2,5 l of the *S. terrestris* inoculum prepared as explained above (Netzer et al.1985). The inoculum of *S. terrestris* in the mix was quantified by the following procedure: 1 g of soil was placed in 100 ml of sterile water and processed according to the serial dilution method and inoculated on PDA (Netzer et al. 1985). In addition, bacterial-inoculated plants were re-inoculated with BsA01 30 days post-transplant in a similar manner as previously described.

### Field experiment

#### Plant material preparation

For this trial, seed bio-priming was performed according to Bisen et al. (2015) and Mahmood et al. (2016). BsA01 was cultured in BHI broth at 28 °C for 24 h. After disinfection with 3% sodium hypochlorite solution, seeds were rinsed and drained for 10 min at room temperature. Half of the seeds were immersed in the *B. subtilis* suspension (OD 1.3). The other half was immersed in BHI broth without bacteria and was used as controls. Seeds suspensions were maintained for 24 h at 28 °C and 170 rpm. Both groups of seeds were dried at room temperature and then sown in the field with a manual seeding machine. In addition, two more inoculations with 1 ml of the same BsA01 suspension (OD=1.3, per plant) were carried out at 60 and 90 days after plant emergence. Plots were irrigated by flooding, as it is normally done in extensive onion crop production systems. Half of the bacteria-inoculated plants and half of the non inoculated plants were inoculated with the pathogen. For this purpose, the St inoculum was multiplied and prepared as indicated before and the pathogen suspension was poured on the planting line, spreading 2 l in 2.5 m. Three replicates of each treatment were carried out in 10 m^2^ subplots.

### Experiment design and variables evaluated

As it was previously stated, for both trials four treatments were defined: **C**: control onion plants, neither inoculated with *B. subtilis* nor *S. terrestris*; **Bs**: plants inoculated only with *B. subtilis*; **St**: plants inoculated only with *S. terrestris*; **Bs-St**: plants inoculated with *B. subtilis* and *S. terrestris*.

In the greenhouse experiment, 16 plants were measured in each evaluation, while in the field experiment 21 plants were measured each time. Symptoms of severity and photosynthetic variables were evaluated in both experiments, while other parameters such as growth and performance were only evaluated in the field experiment.

#### Disease severity and biocontrol assessment

Pink root severity was assessed every 30 days using the scale proposed by Piccolo and Galmarini (1994) with modifications. The scale proposes 6 severity levels (0-5) and classifies infected roots according to the percentage of roots with pink discoloration and or flattened roots (0 = healthy plant or plant without symptoms, 1 = plant affected by 1-20 %, 2 = 21-40%, 3 = 41-60%, 4 = 61-80%, 5 = 81-100%). To determine this parameter, plants of each treatment were uprooted and washed, and the percentage of affected roots was calculated. These values were used to build the disease progress curve. The area under the disease progress curve (AUDPC) was calculated according to Campbell and Madden, 1990.

Finally, to measure the biocontrol efficacy of BsA01 we applied the formula proposed by Kakar et al. (2018) as follow:

Biocontrol efficacy = [(CS-TS) / CS] × 100, where CS refers to the severity of treatment St and TS refers to the severity of the treatment Bs + St.

Values used for calculation were obtained at 130 days after sowing for the greenhouse trial and at 100 days for the field trial.

#### Growth parameters

As growth parameters we evaluated: dry matter, length of leaves and roots. In addition, the diameter of the bulbs and yield were also estimated since both are important for commercial qualification of onion. These variables were measured only in the field trial at 40, 70 and 100 days after sowing of the crop.

#### Photosynthetic variables

Net CO_2_ assimilation rate (*A*) and stomatal conductance (*gs*) were measured using a portable photosynthesis system (LICOR 6400XT, Lincoln, NA, USA). All readings were taken between 11 am and 2 pm, on the youngest, fully expanded leaves at 70, 100 and 130 days. The light inside the cuvette was set at saturating photosynthetic photon flux density (PPFD) of 1500 µmol m^-2^ s^-1^. The temperature of measurement was fixed at 25°C and the CO_2_ reference was kept at 400 µmol m^-2^ s ^-1^.

Afterwards, gas exchange (GS) measurements and fast chlorophyll A fluorescence (ChlF) emission were measured using a Pocket PEA (Plant Efficiency Analyser, Hansatech Instruments Ltd., King’s Lynn, Norfolk, UK). The measurements were performed on the middle part of adaxial leaf blades, as described for GS parameter, after a dark adaptation of (> 25 min) using light-exclusion clips. Upon 3s light exposition at light intensity of 3500 mmol photons m ^-2^ s ^-1^ and peak wavelength of 650 nm, the fluorescence signal was collected at a maximum frequency of 105 points s-1 (each 10 μs). The basic parameters derived from the ChlF transient data were used to perform the OJIP test as described by Streasser and co-workers. The OJIP test detects real-time variations of the photosynthetic apparatus and plant vitality in physiological or stress conditions (Streasser et al. 2004). These basic parameters are minimum fluorescence emission recorded at 20 µs (F_0_ or O step); fluorescence intensities at 2 ms and 3 ms (J and I step respectively) and maximum fluorescence, P step ∼ 300 ms (F_m_).

Under open field trial, simultaneous measurements of net CO2 assimilation rate, stomatal conductance, as well as modulated ChlF emission were recorded using the same equipment (LICOR 6400XT) described in the previous paragraph. The measurements were done with the same criteria used under greenhouse experiment at day 90 after sowing of the crop. The parameters related to the PSII electron transport activity were calculated as described previously (Baker et al., 2008): PSII maximum quantum efficiency (Fv’/Fm’ = Fm’ - F0 / F0’); PSII operating efficiency (Fq’ / Fm’ = Fm’ – Fs’ /Fm’), where Fs is steady-state fluorescence.

### Statistical analysis

To analyze severity, samples were randomly extracted within each treatment and according to the categorical scale previously stated. The variable was transformed to binomial in order to perform this type of analysis. Therefore, percentages ≤ 40% (categories of scale 0, 1, 2 and 3) of root affected by St were considered as 0, while percentages > 40% were included in values of 1. We then applied generalized linear models, logit link (logistic regression) with the statistical program SPSS (IBM Corp.). In both trials, two-way ANOVA was used to evaluate the differences among physiological variables. The analyses were conducted using INFOSTAT.®, version 2017. Means comparison were performed using post-hoc Fisher’s (P≤0.05) test.

In the field experiment, ANOVA of repeated measures was applied to examine the quantitative parameters, specifying the plot as the repeated term and using the linear procedure of mixed models in INFOSTAT® 2017 (Di Rienzo et al. 2017). We captured the variation among the plots directly by modeling the variance-covariance matrix of the residuals on each task for each plot. The best fitting covariance structure for the residuals across trials was autoregressive (AR1) that resulted in improved fit of the model. The multiple comparisons between the means of the treatments were determined by using Fisher test (P≤0.05) and the measurement of the variability reported with the mean corresponds to the standard error.

## Results

In both trials, all plants inoculated only with *S. terrestris* showed pink root symptoms (Fig. 1) indicating that the infectious inoculum was effective and plants were susceptible to the pathogen.

**Fig 1.**
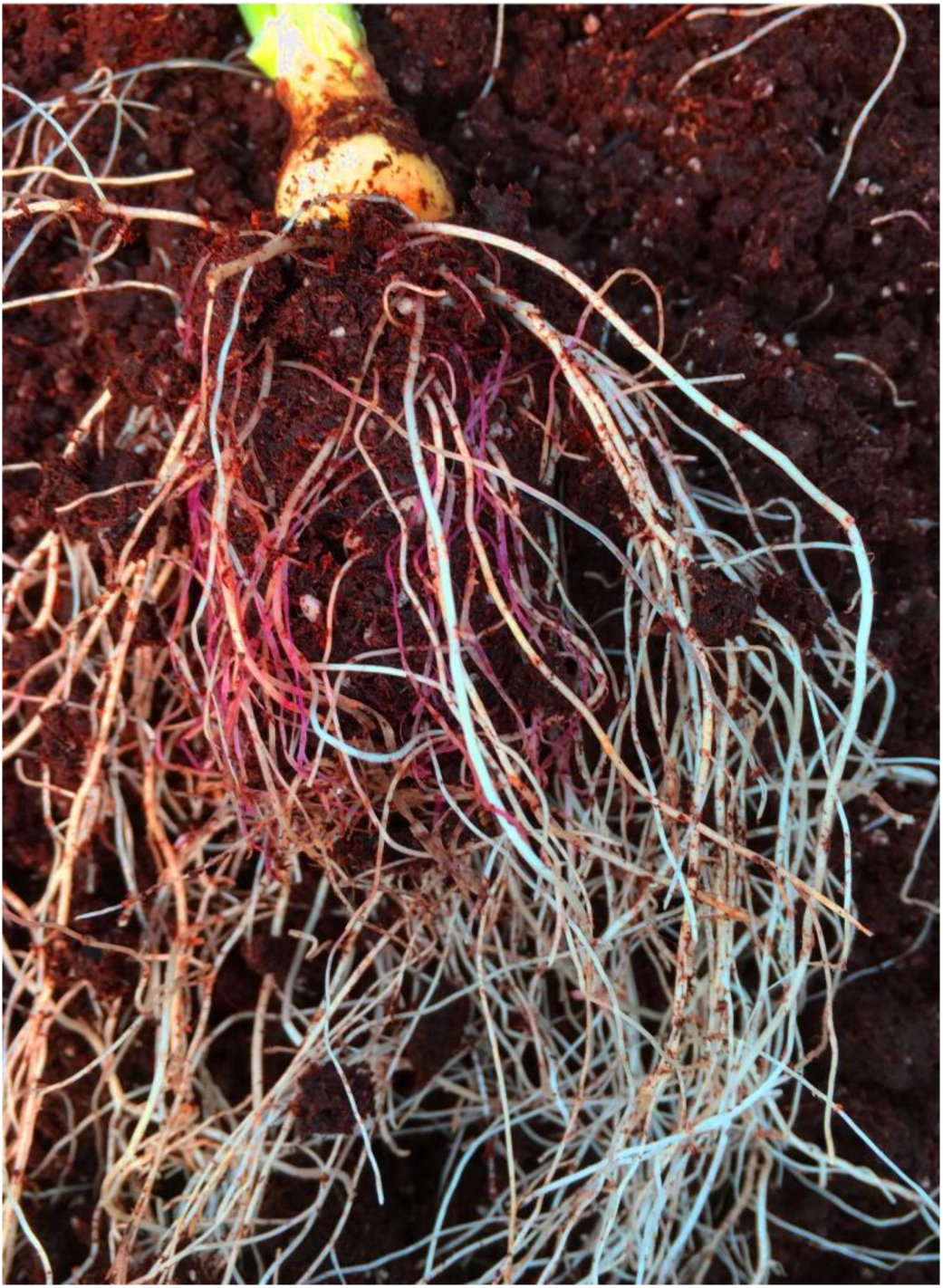
Pink root disease symptoms in onions caused by the soil-borne fungal pathogen *Setophoma terrestris*. Typical symptomatology is observed in the cultivation of onions, a pink to purple coloration of the roots.

### Pink root severity and biocontrol under greenhouse conditions

The progress of the disease was evaluated by the calculation of AUDPC (Fig 2a). For treatment St the AUDPC was 187.5 while for treatment Bs-St was 95.6. The disease severity was significantly reduced when the bacterial strain was applied. We found by logistic regression analysis that Bs-St treatment showed 7 times less probability of disease occurrence (Exp (B) = 7.00, *P* = 0.005 and Interval of Confidence 95% = 1.822-26.887). According to the average severity values obtained from measurements at day 130, the biocontrol efficiency was of 50.9%.

**Fig 2.**
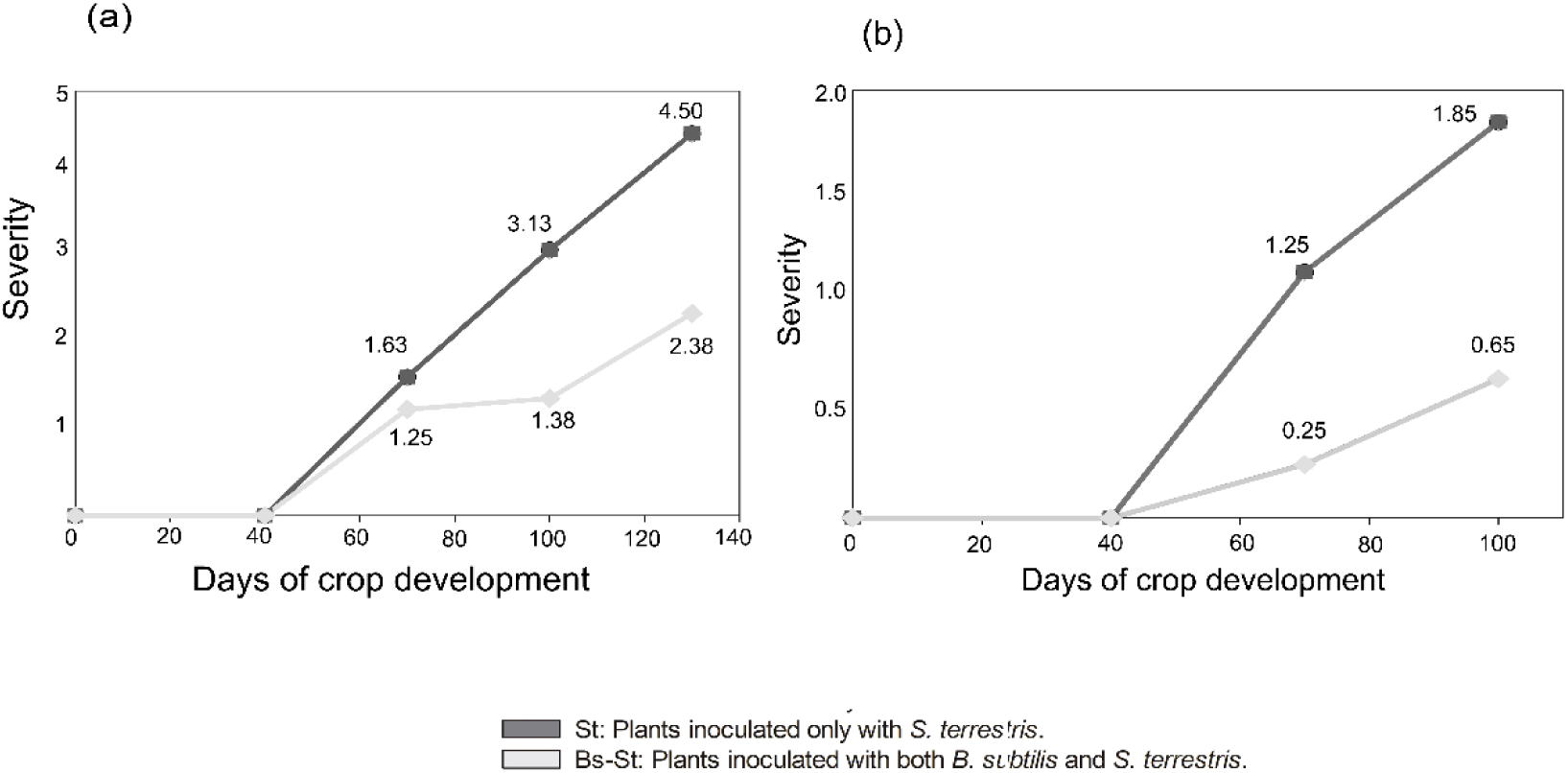
Area under the Pink root disease progress curve (AUDPC) for onion seedlings inoculated with St = *Setophoma terrestris*, St-Bs = *Setophoma terrestris* + *Bacillus subtilis* ALBA01 under greenhouse (a) and field conditions (b).

### Pink root and biocontrol effects on onion plant growth under field conditions

The severity at day 100 for St treatment was 1.85, with damage not only observed at the root level but also in the aerial part of the plants, with necrosis at the tips of the leaves. Treatment Bs-St resulted in a final severity value of 0.65 (Fig. 2b). A logistic regression was performed to determine the severity of both treatments. The logistic regression model was statistically significant, (Wald= 5.112; *P* =0.024). The model explained 27.0% (Nagelkerke R2) of the variance in the disease and correctly classified 77.5% of the cases. Plants inoculated with BsA01 were 12.67 times less likely to have disease than control plants (Odd Ratio = 12.667, 95% CI = 1.402 - 114.419, p = 0.024).

The AUDPC value of treatment St was 72.75, while treatment Bs-St presented a value of 17.25 (Fig. 2b). The biocontrol efficiency obtained reached a value of 64.86 %.

### Quantitative plant growth parameters in the field experiment

We found significant differences in dry matter measurements, with St treatment showing a 67.45% reduction in dry matter compared to control plants (F= 5.28; P = 0.0015, Fig. 3a). Bs-St plants showed an increase in leaf length of 11.36% compared to control treatment, while the lowest value was observed for the treatment with St, which showed exhaustion of 7.67% also compared to control (F= 17.06; P = 0.01, Fig. 3b).

**Fig 3.**
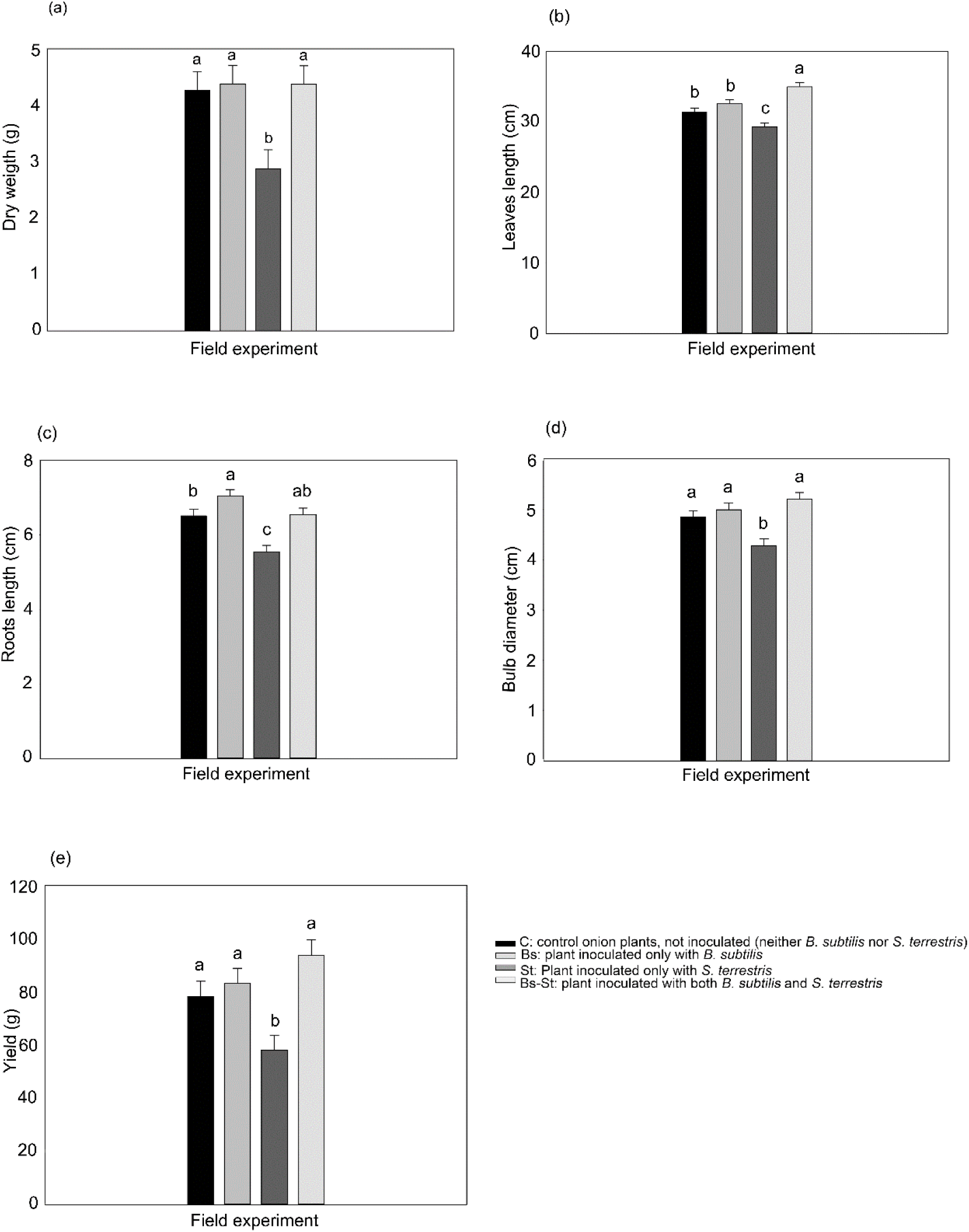
Effects on the quantitative parameters of plant growth under greenhouse and field conditions. Dry weight (a); leaf length of the leaves (b); length of the root (c); diameter of the bulb (d); performance (e) were mediated in the treatments: C; Bs; St; St-Bs The measurements were made every 30 days in the respective environmental conditions. The data are means ± SE and different letters show significant differences with a value of p < 0.05.

Inoculation with BsA01 determined significant differences in root length. Treatment Bs showed higher root length compared to St and control plants but they do not present significant differences with Bs-St treatment. (F = 12.39, P <0.0001, Fig. 3c). BsA01 inoculation resulted in higher bulb diameter and yield, since plants of Bs-St, Bs and control treatments were significantly different from the St treatment (bulb diameter: F=9.08; P = <0.0001; yield: F=6. 85; P = 0.0004, Fig. 3d-3e).

### Gas exchange and ChlF analysis in plants grown under greenhouse conditions

Thirty days after inoculation, both net CO_2_ assimilation rate and stomatal conductance (Fig. 4a -b), were significantly reduced in those plants infected by *S. terrestris.* The alleviating effect of BsA01 on the pathogen deleterious effect, (treatment Bs-St) was only partially and the negative effect was apparent when net photosynthesis and stomatal conductance were measured.

**Fig 4.**
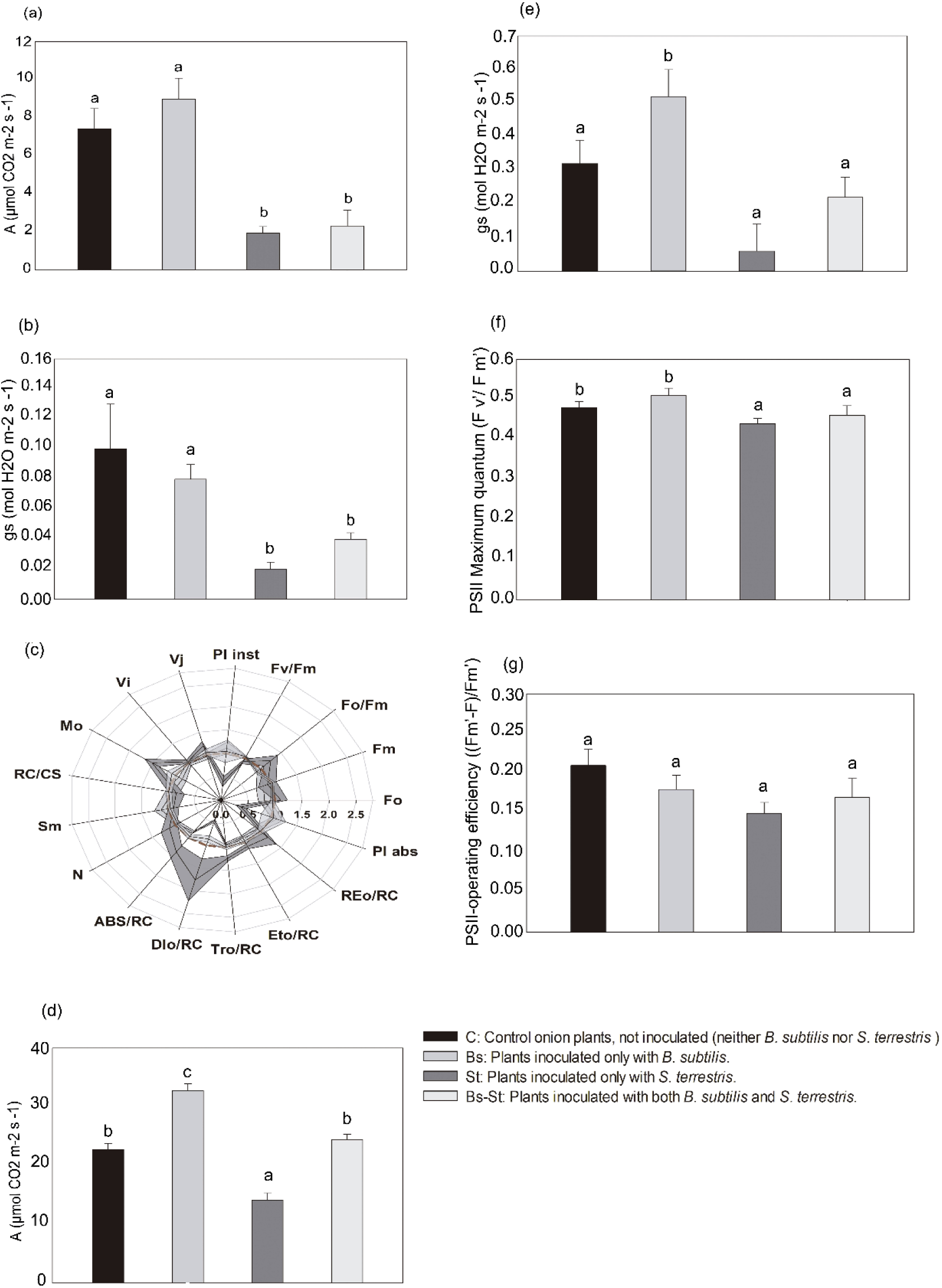
Net rate of assimilation of CO2 (a) and stomatal conductance (b), in onion plants of the treatments: C; Bs; St; St-Bs, measured 70 days after sowing under greenhouse conditions. The data are means ± SE and different letters show significant differences with a value of p <0, 05. Radar plot showing the functioning of the PSII by a set of OJIP-test parameters measured the day 70 after sowing in onion plants (C; Bs; St; St-Bs). The values were normalized to the control plants. For the description and calculation of this parameters see Supplementary table1. Net CO2 assimilation rate (d), stomatal conductance (e), PSII maximum quantum efficiency (f), and PSII operating efficiency (g). Data are means ± SE and different letters show significant differences with a p-value < 0, 05.

Measurements of ChlF provide a non-invasive means of probing the dynamics of photosynthesis. The relative values of the OJIP parameters (relative to control), were plotted on a radar plot in order to get an overall picture of the incidence of *S. terrestris* and the PGPR effect of BsA01 on the light reaction of photosynthesis (Fig. 4c).

*S. terrestris*-infected plants (St) showed an abrupt fall (∼50%) of the maximum quantum yield. Moreover, the dissipative light energy mechanism per reaction center parameter sharply increased (by more than two folds) in St plants, indicating that those plants needed to activate the mechanism of dissipation as heat in the Photosystem II (PSII) antenna, which commonly occurs under stressful conditions. It is interesting to note that control plants (C), Bs and Bs-St showed neither signs of inactivation of reaction centers (changes in F_v_/F_m_) nor experienced an increment in the DIo/RC (Fig 4c). In addition, there was an imbalance in the energy flow parameters such as light energy absorption (ABS/RC), electron trapping (TRo/RC), linear electron transport (ETo/RC) and the electron flow at the PSI acceptor side (REo/RC) by the presence of the pathogen, while such parameters were unresponsive in the rest of the treatments (Fig. 4c). The index of vitality of the infected plants was drastically reduced as indicated by the absolute (P_abs_) and instantaneous performance index (P_inst_), while in plants belonging to C, Bs and Bs-St treatments no significant changes were recorded (Fig. 4c). Finally, *S. terrestris* also affected the initial slope of the fluorescence transient (Mo) parameter, which express the rate of reaction center (RC) closure and the variable fluorescence emission at the J step (Vj), while in the *Bacillus*-inoculated (Bs-St) and control plants (C) no significant changes were registered (Fig. 4c).

### Gas exchange and ChlF analysis in plants grown under field conditions

Under field conditions, the pernicious effect of *S. terrestris* on CO2 assimilation resembled those observed in the greenhouse trial. In infected plants (St), net CO2 assimilation rate was greatly inhibited (∼100% in relation to C plants) (Fig. 4d). Such response was correlated to the stomatal conductance, which showed a similar trend (Fig. 4e). On the other hand, net photosynthesis of the plants inoculated with BsA01 was significantly higher than those of control plants.

The PSII maximum quantum efficiency assessed by simultaneous measurements of ChlF showed a similar pattern to those observed in photosynthetic evaluations (A and gs), but the intensity at which it dropped was slighter (Fig. 4f). On the other hand, none of the treatments showed signs of alteration on the operating PSII efficiency (Fig. 4g).

## Discussion

It has been extensively reported that some soil microbes, such as fungi and bacteria, can profusely colonize the rhizosphere where the biology and chemistry of the soil are highly influenced by root exudates (Lareen et al. 2016; Khalid et al. 2009). Among the rhizospheric inhabitants and their outstanding activities, PGPR provide a front defence line against virus, bacterial and fungal attack (Glick 1995; Kloepper et al. 1989; Bhattacharyya et al. 2012).

In a previous study, the biocontrol ability of BsA01 against *S. terrestris* was tested in dual cultures *in vitro* (Albarracín Orio et al. 2016a), but it has never been challenged to the pathogen *in vivo*. Here, we aimed to test the potential biocontrol and plant growth promoting traits of BsA01 under greenhouse and fields conditions.

Our findings suggest that *Bacillus subtilis* ALBA01 exhibits a significant effect of bioprotection against the soil-borne pathogen *S. terrestris in vivo* by reducing the damage caused by *S. terrestris* through maintaining the photosynthesis activity and stimulating plant growth. The average biocontrol efficiency of BsA01 was of 57.42%. These results were in agreement with the AUDPC values that revealed a high final severity in plants of St treatment. Thus, treatments infected with the pathogen, St and St-Bs, showed clear symptoms of pink root disease, however, they were clearly mitigated by the presence of BsA01. Moreover. we found a lower probability of disease occurrence in St-Bs. BsA01 treatment resulted in a 35% higher levels of the dry matter compared to St treatment. A similar pattern was observed in other biometric parameters such as leaf and root length, diameter of the bulb and yield. These results might indicate a high probability that the *S. terrestris*-infected plants inoculated with BsA01did not become diseased and had a normal development.

These positive and evident effects could be based on multiple causes because the biological control mechanisms of *B. subtilis* include the production of antifungal substances, the induction of plant resistance (Kloepper 2004) and the promotion of plant growth (Wang et al. 2018). These characteristics of BsA01 can be attributed because the plants of the Bs-St treatments did not get diseased or had less damage. In addition, this effect of BsA01 could be due to an appropriate time of stimulation or to the colonization of the root by the PGPR triggering the process of systemic induced resistance (Timmusk et al. 1999), where the plant exhibits an increase in the level of resistance to infection by a pathogen by activating hormone and sugar signals (Shoresh et al. 2010). This cascade of signals might explain the effects on the opening of the stomatal and, consequently, on the net rate of assimilation of CO2. From previous studies, it is known that abscisic acid synthesis mediates the closure of the stomatal as a primary defense mechanism (Bauer et al. 2013; McAdam et al. 2016). In both trials, we observed that plants infected with *S. terrestris* markedly reduced the net assimilation rate of CO2. However, in the Bs-St treatments, the negative effect of the pathogen *S. terrestris* on the photosynthetic rate was notoriously attenuated, demonstrating its efficacy as BCA.

On the other hand, the functional alteration of the photosynthetic apparatus related to pathogens is possible (Bauriegel et al. 2014; Berger et al. 2007). In the other phase of the photosynthetic process (reaction to light in PSII), *S. terrestris* infection induced changes in most of the OJIP parameters. Infected plants showed signs of photoinhibition, denoted by the classical parameter Fv / Fm or “maximum quantum yield”. This result indicates that *S. terrestris* induced the irreversible closure of some PSII (RC) reaction centers, which caused the increase of the minimum fluorescence emission (Fo) (Franklin et al. 1992; Yamane et al. 2000). This can generate over-excitation of the reaction centers of photosystem II (PSII) and the formation of reactive oxygen species (ROS). To avoid over-excitation of the PSII and damage to the photosynthetic apparatus, the energy that does not take the photochemical path can be dissipated mainly as heat, as observed by the activation of the dissipative light energy mechanism. The absolute yield index (PIabs), a highly sensitive indicator of the vitality of the plant, also decreased in plants infected with *S. terrestris*. However, in control and St-Bs treatments, PIabs remained unchanged and showed similar values among them. These results are consistent with that reported by Ajigboye et al. (Ajigboye et al. 2016) and Zhori et al. (Zhori et al. 2015), who showed the incidence of various pathosystems on different plant species using the OJIP analysis. A similar trend was observed in the field trial. The maximum quantum efficiency of PSII decreased by the effect of infection by St compared to control and St-Bs treatments. In this case, the elevation of Fv ‘/ Fm’ occurred due to the effect of BsA01, which correlated positively with a significant increase in stomatal conductance and, consequently, a higher photosynthetic rate.

The effect on root growth promotion that we observed for BsA01 inoculated plants is in agreement with results shown by Bouizgarne, B. (Bouizgarne 2013) where *B. subtilis* triggered morphogenetic changes in the *Arabidopsis thaliana* root system, based on the stimulation of the primary root and/or the lateral root development. The continuous generation of new roots in addition to the BsA01 fungal inhibition capacity (Albarracín Orio et al. 2016a) could be responsible for the normal development of the onion plants. Our results clearly show that *B. subtilis* ALBA01 mitigate pink root damage, proving that it can be used as an effective biocontrol agent against *S. terrestris*.

## Supporting information

Formulas and glossary of terms used by the OJIP-test

## References

Ajigboye OO, Bousquet L, Murchie EH, Ray RV (2016). Chlorophyll fluorescence parameters allow the rapid detection and differentiation of plant responses in three different wheat pathosystems. Functional plant biology 43(4), 356–369. doi:10.1071/FP15280

Albarracín Orio, A.G., Brücher, E. & Ducasse, D.A. (2016a) A strain of *Bacillus subtilis subsp. subtilis* shows a specific antagonistic activity against the soil-borne pathogen of onion *Setophoma terrestris*. Eur J of Plant Pathol 144:217–223. doi:10.1007/s10658-015-0762-0

Albarracín Orio AG, Tobares RA, Ducasse DA, Smania AM. (2016b). Draft genome sequence of Bacillus subtilis ALBA01, a strain with antagonistic activity against the soilborne fungal pathogen of onion *Setophoma terrestris.* Genome Announc 4(3):e00455–16. doi:10.1128/genomeA.00455-16.

Baker NR (2008) Chlorophyll fluorescence: a probe of photosynthesis *in vivo.* Annu Rev Plant Biol 59:89–113. doi:10.1146/annurev.arplant.59.032607.092759

Bauer H, Ache P, Lautner S, et al (2013) The stomatal response to reduced relative humidity requires guard cell-autonomous ABA synthesis. Curr Biol 23:53–57. doi:10.1016/j.cub.2012.11.022

Bauriegel E, Herppich W (2014) Hyperspectral and Chlorophyll Fluorescence Imaging for Early Detection of Plant Diseases, with Special Reference to *Fusarium* spec. Infections on Wheat. Agriculture 4:32–57. doi:10.3390/agriculture4010032

Berger S, Benediktyová Z, Matouš K, et al (2007) Visualization of dynamics of plant-pathogen interaction by novel combination of chlorophyll fluorescence imaging and statistical analysis: Differential effects of virulent and avirulent strains of *P. syringae* and of oxylipins on *A. thaliana*. J Exp Bot 58:797–806. doi:10.1093/jxb/erl208

Berger S, Papadopoulos M, Schreiber U, Kaiser W, Roitsch T (2004) Complex regulation of gene expression, photosynthesis and sugar levels by pathogen infection in tomato. Physiol Plant 122:419–428. doi:10.1111/j.1399-3054.2004.00433.x

Bhattacharyya PN, Jha DK (2012) Plant growth-promoting rhizobacteria (PGPR):Emergence in agriculture. World J Microbiol Biotechnol 28:1327–1350. doi:10.1007/s11274-011-0979-9

Bisen K, Keswani C, Mishra S, Saxena A, Rakshit A, Singh HB (2015). Unrealized potential of seed biopriming for versatile agriculture. In Nutrient Use Efficiency: from Basics to Advances 193–206. doi:10.1007/978-81-322-2169-2_13

Bouizgarne B (2013) Bacteria for plant growth promotion and disease management. In: Bacteria in agrobiology: disease management. Springer 15–47. doi:10.1007/978-3-642-33639-3_2

Cai, X. C., Liu, C. H., Wang, B. T., & Xue, Y. R. (2017). Genomic and metabolic traits endow *Bacillus velezensis* CC09 with a potential biocontrol agent in control of wheat powdery mildew disease. Microbiological research, 196:89–94. doi:10.1016/j.micres.2016.12.007

Campbell CL, Madden L V (1990) Introduction to plant disease epidemiology. John Wiley & Sons.

Costa Pinto, L.S., Azevedo, J.L., Pereira, J.O., Carneiro Vieira, M.L., Labate, C.A., (2008). Symptomless infection of banana and maize by endophytic fungi impairs photosynthetic efficiency. New Phytol. 147, 609–615. doi:10.1046/j.1469-8137.2000.00722.x

de Gruyter J, Woudenberg JHC, Aveskamp MM, et al (2010) Systematic reappraisal of species in *Phoma* section *Paraphoma, Pyrenochaeta* and *Pleurophoma*. Mycologia 102:1066–1081. doi:10.3852/09-240

Di Rienzo J.A., Casanoves F., Balzarini M.G., Gonzalez L., Tablada M., Robledo C.W. InfoStat versión 2017. Grupo InfoStat, FCA, Universidad Nacional de Córdoba, Argentina. URL http://www.infostat.com.ar

Esfahani, M. N., & Pour, B. A. (2008). Differences in resistance in onion cultivars to pink root rot disease in Iran. Journal of general plant pathology, 74(1), 46–52.,

Franklin LA, Levavasseur G, Osmond CB, et al (1992) Two components of onset and recovery during photoinhibition of *Ulva rotundata*. Planta 186:399–408. doi:10.1007/BF00195321

Glick BR (1995) The enhancement of plant growth by free-living bacteria. Can J Microbiol 41:109–117. doi:10.1139/m95-015

Hansatech I (2006) Operations manual: Handy PEA, Pocket PEA y PEA Plus

Hansen, H.N. (1929). Etiology of the pink-root disease of onions. Phytopathology 19:691–704

Kakar KU, Nawaz Z, Cui Z, et al (2018) Rhizosphere-associated *Alcaligenes* and *Bacillus* strains that induce resistance against blast and sheath blight diseases, enhance plant growth and improve mineral content in rice. J Appl Microbiol 124(3), 779–796. doi:10.1111/jam.13678

Khalid A, Arshad M, Shaharoona B, Mahmood T (2009) Plant growth promoting rhizobacteria and sustainable agriculture. In: Microbial Strategies for Crop Improvement Springer 133–160. doi:10.1007/978-3-642-01979-1_7

Kloepper JW, Lifshitz R, Zablotowicz RM (1989) Free-living bacterial inocula for enhancing crop productivity. Trends Biotechnol 7:39–44. doi:10.1016/0167-7799(89)90057-7

Kloepper JW, Ryu C, Zhang S (2004) Induced Systemic Resistance and Promotion of Plant Growth by *Bacillus* spp. Phytopathology 94:1259–1266. doi:10.1094/PHYTO.2004.94.11.1259

Kloepper JW, Schroth MN (1978) Plant growth-promoting rhizobacteria on radishes. In Proc. of the 4th Internet. Conf. on Plant Pathogenic Bacter, Station de Pathologie Vegetale et Phytobacteriologie 2: 879–882.

Lafi J. G. (2011) Caracterización morfológica, fisiológica, patogénica y molecular de aislados argentinos de Phoma terrestris Hansen. (tesis de postgrado). Universidad Nacional de Cuyo, Mendoza.

Lareen A, Burton F, Schäfer P (2016) Plant root-microbe communication in shaping root microbiomes. Plant Mol Biol 90:575–587. doi:10.1007/s11103-015-0417-8

Leal N, Lastra R (1984) Altered metabolism of tomato plants infected with tomato yellow mosaic virus. Physiol Plant Pathol 24:1–7. doi:10.1016/0048-4059(84)90067-5

Lee C-J, Lee J-T, Moon J-S, et al (2007) Effects of solar heating for control of Pink Root and other soil-borne diseases of onions. Plant Pathol J 23:295–299. doi:10.5423/PPJ.2007.23.4.295

Linardelli C., Lafi, J., Puglia, C., Tarquini, A., Soto, A., Echevarría, S. (2008a) Comportamiento a campo de cuatro aislados de *Phoma terrestris* sobre dos cultivares de cebolla. Marinelli A. D. (Presidencia). 1° Congreso Argentino de Fitopatología. Córdoba, Argentina. ISBN:978-987-24373-0-51

Linardelli, C., Tarquini, A., Lafi, J., Echevarria, S. (2008b) Variabilidad patogénica de aislados de *Phoma terrestris* presentes en argentina. Marinelli A. D. (Presidencia). 1° Congreso Argentino de Fitopatología. Córdoba, Argentina. ISBN:978-987-24373-0-51

Mahmood A, Turgay OC, Farooq M, Hayat R (2016). Seed biopriming with plant growth promoting rhizobacteria: a review. FEMS Microbiol Ecol, 92(8). doi:10.1093/femsec/fiw112

Marzu, J. C., Straley, E., & Havey, M. J. (2018). Genetic Analyses and Mapping of Pink-Root Resistance in Onion. Journal of the American Society for Horticultural Science, 143(6), 503–507

McAdam SAM, Manzi M, Ross JJ, et al (2016) Uprooting an abscisic acid paradigm: Shoots are the primary source. Plant Signal Behav 11:652–659. doi:10.1080/15592324.2016.1169359

Netzer D, Rabinowitch HD, Weintal C (1985) Greenhouse technique to evaluate onion resistance to pink root. Euphytica 34:385–391. doi:10.1007/BF00022933

Phookamsak R, Liu JK, Manamgoda DS, Chukeatirote E, Mortimer PE, Mckenzie EH, Hyde KD (2014). The sexual state of Setophoma. Phytotaxa, 176 (1), 260–269

Piccolo, R. J. and Galimarini, C. R. 1994. Epidemiological characteristics of onion pink root (*Phoma terrestris* (Hansen)) resistance. RIA, Revista-de-Investigaciones Agropecuarias 25: 1–13.

Porter, I. J., Merriman, P. R., & Keane, P. J. (1989). Integrated control of pink root (Pyrenochaeta terrestris) of onions by dazomet and soil solarization. Australian Journal of Agricultural Research, 40 (4), 861–869.

Rinland ME, Gómez MA (2015) Isolation and characterization of onion degrading bacteria from onion waste produced in South Buenos Aires province, Argentina. World J Microbiol Biotechnol 31:487–497. doi:10.1007/s11274-015-1803-8

Roitsch T, Balibrea ME, Hofmann M, et al (2003) Extracellular invertase: key metabolic enzyme and PR protein. J Exp Bot 54:513–524. doi:10.1093/jxb/erg050

Sajitha KL, Dev SA, Maria Florence EJ (2018) Biocontrol potential of *Bacillus subtilis* B1 against sapstain fungus in rubber wood. Eur J Plant Pathol 150: 237–244. doi:10.1007/s10658-017-1272-z

Sawinski K, Mersmann S, Robatzek S, Böhmer M (2013) Guarding the green: pathways to stomatal immunity. Mol Plant-Microbe Interact 26:626–632. doi:10.1094/MPMI-12-12-0288-CR

Schwartz HF, Mohan SK (2007) Compendium of onion and garlic diseases and pests. APS Press St Paul, MN 1008:127. doi:10.1094/9780890545003.002

SENASA. Servicio Nacional de Sanidad y Calidad Agroalimentaria (2014). La cebolla, embajadora de la calidad hortícola argentina. http://www.senasa.gob.ar/senasa-comunica/infografias/la-cebolla-embajadora-de-la-calidad-horticola-argentina, consulta:_abril 2017.

Shoresh M, Harman GE, Mastouri F (2010) Induced Systemic Resistance and Plant Responses to Fungal Biocontrol Agents. Annu Rev Phytopathol 48:21–43. doi:10.1146/annurev-phyto-073009-114450

Singh A, Gupta R, Pandey R (2016) Rice seed priming with picomolar rutin enhances rhizospheric *Bacillus subtilis* CIM colonization and plant growth. PLoS One 11:e0146013. doi:10.1371/journal.pone.0146013

Timmusk, S., & Wagner, E. G. H. (1999). The plant-growth-promoting rhizobacterium Paenibacillus polymyxa induces changes in Arabidopsis thaliana gene expression: a possible connection between biotic and abiotic stress responses. Molecular plant-microbe interactions, 12 (11), 951–959.

Wang, X. Q., Zhao, D. L., Shen, L. L., Jing, C. L., & Zhang, C. S. (2018). Application and Mechanisms of Bacillus subtilis in Biological Control of Plant. Role of Rhizospheric Microbes in Soil: Volume 1: Stress Management and Agricultural Sustainability, 225.

Xiang N, Lawrence KS, Kloepper JW, Donald PA, McInroy JA (2017) Biological control of *Heterodera glycines* by spore-forming plant growth-promoting rhizobacteria (PGPR) on soybean. PLoS One 12(7): e0181201. doi:10.1371/journal.pone.0181201

Yamane Y, Shikanai T, Kashino Y, et al (2000) Reduction of Q (A) in the dark: Another cause of fluorescence F(o) increases by high temperatures in higher plants. Photosynth Res 63:23–34. doi:10.1023/A:1006350706802

Zhori A, Meco M, Brandl H, Bachofen R (2015) In situ chlorophyll fluorescence kinetics as a tool to quantify effects on photosynthesis in *Euphorbia cyparissias* by a parasitic infection of the rust fungus *Uromyces pisi.* BMC Res Notes 8:1–7. doi:10.1186/s13104-015-1681-z

