## Supplementary material for "*Bacillus subtilis* ALBA01 can mitigate onion pink root symptoms caused by *Setophoma terrestris*": Formulas and glossary of terms used by the OJIP-test

**Supplementary Table**. Formulas and glossary of terms used by the OJIP-test (fig.4c).

| | |
|---|---|
| Fo | Estimated emission by excited chlorophyll a in PSII reaction centers (RCs) after dark adaptation, when the first stable electron acceptor in PSII ($Q_A$) is fully oxidized. |
| Fm | Maximum fluorescence obtained at saturating light intensity and when $Q_A$ is fully reduced. |
| Fv | Maximum capacity for photochemical quenching. Calculated as Fv=Fm-Fo |
| Fv/Fm | Maximum quantum efficiency of PSII. Its values decrease in stressed plant samples. |
| Fv/Fo | Size and number of active reaction centers RC. Reflect impairment and downregulation of PSII photochemistry and low electron transport. |
| Vj | Relative variable fluorescence at 2 ms, step J. Calculated as Vj=(FJ – F0)/(Fm – F0). |
| Vi | Relative variable fluorescence at 30 ms, step I. Calculated as Vi=(Fi – F0)/(Fm – F0). |
| Sm | Measure of the energy needed to close all reaction centers. |
| N | Qa turnover number, it indicates how many times Qa has been reduced in the time span from 0 to $t_{Fmax}$ . |
| Area | Total complementary area between fluorescence induction curve and F = Fm |
| Dlo/RC | Ratio of total dissipation to the amount of active RCs. It increases due to the high dissipation of the inactive RCs. |
| TRo/RC | Trapped energy flux per RC (at t=0). |
| ERo/RC | Electron transport flux per RC (at t=0) |
| REo/RC | Electron flux reducing end electron acceptors at the PSI acceptor side, per RC |
| $PI_{abs}$ | Performance index on absorption basis, it is an indicator of sample vitality. ABS is flux of photons absorbed by the antenna pigments Chl |
| RC/CS | Denseity of reaction centers (RC) per corss section (CS) |
